# Autophosphorylation of the KaiC-like protein ArlH inhibits oligomerisation and interaction with ArlI, the motor ATPase of the archaellum

**DOI:** 10.1101/2021.03.19.436134

**Authors:** J. Nuno de Sousa Machado, Leonie Vollmar, Julia Schimpf, Paushali Chaudhury, Rashmi Kumariya, Chris van der Does, Thorsten Hugel, Sonja-Verena Albers

## Abstract

Motile archaea are propelled by the archaellum, whose motor complex consists of the membrane protein ArlJ, the ATPase ArlI, and the ATP-binding protein ArlH. Despite its essential function and the existence of structural and biochemical data on ArlH, the role of ArlH in archaellum assembly and function remains elusive. ArlH is a structural homolog of KaiC, the central component of the cyanobacterial circadian clock. Similar to KaiC, ArlH exhibits autophosphorylation activity, which was observed for both ArlH of the euryarchaeon *Pyrococcus furiosus (Pf*ArlH) and the crenarchaeon *Sulfolobus acidocaldarius* (*Sa*ArlH). Using a combination of single molecule fluorescence measurements and biochemical assays, it is shown that autophosphorylation of ArlH is closely linked to the oligomeric state of ArlH bound to ArlI. These experiments also strongly suggest that ArlH is a hexamer in its functional ArlI bound state. Mutagenesis of the putative catalytic residue Glu-57 in *Sa*ArlH results in a reduced autophosphorylation activity and abolished archaellation and motility, suggesting that optimum phosphorylation activity of ArlH is essential for both archaellation and motility.

## Introduction

The ability to escape stressful conditions towards environments where growth is optimal can be found across all domains of life (1). In Archaea, motility can be achieved by passive means, e.g., gas vesicles for floating (2), or by active propulsion through liquid media powered by archaella (3). Archaella are functionally analogous to the bacterial flagellum, but these two nanomachines are structurally distinct and evolved independently (1, 3–5). The archaellum is evolutionarily related to type IV pili (T4P) of bacteria and archaea and to type II secretion systems (T2SS) and shares some biochemical properties with these machineries, e.g., its assembly and rotation is ATP-driven and the pilin and archaellin precursors are processed in a similar manner (7–9). The functional similarities between the flagellum and the archaellum and the repurposing of a T4P with a rotary motor are extraordinary examples of convergent and divergent evolution (3).

The archaellum is a relatively simple motility structure, with the archaellum from the model organism *Sulfolobus acidocaldarius* requiring only seven different structural proteins for its function and a membrane-embedded signal peptidase to cleave the signal peptide of pre-archaellinds (See Fig. 1) (10). These proteins are encoded in the *arl* (**<u>ar</u>**chae**l**lin-related genes) cluster, which was previously named the *fla* cluster (11). The *arl* cluster encodes the structural unit (ArlB, archaellin) of the filament and the ArlX/G/F/H/I/J proteins which constitute the archaellum assembly and rotary machinery (See Fig. 1) (4). ArlX is found exclusively in Crenarchaeotes and its function was proposed to be replaced in Euryarchaeotes by the ArlC/D/E proteins (Fig. 1) (3, 12). It was proposed that this is an adaptation to the bacterial-like chemotaxis system present in Euryarchaeotes (13–16). In *Haloferax volcanii*, ArlC/D/E localise to the cell poles in an ArlH-dependent manner, where they are the direct receivers of chemotactic signals transduced by CheF (17, 18). These cytosolic proteins have further been hypothesized to interact with the polar cap that is present in Euryarchaea (See Fig. 1) (18–21).

**Figure 1.**
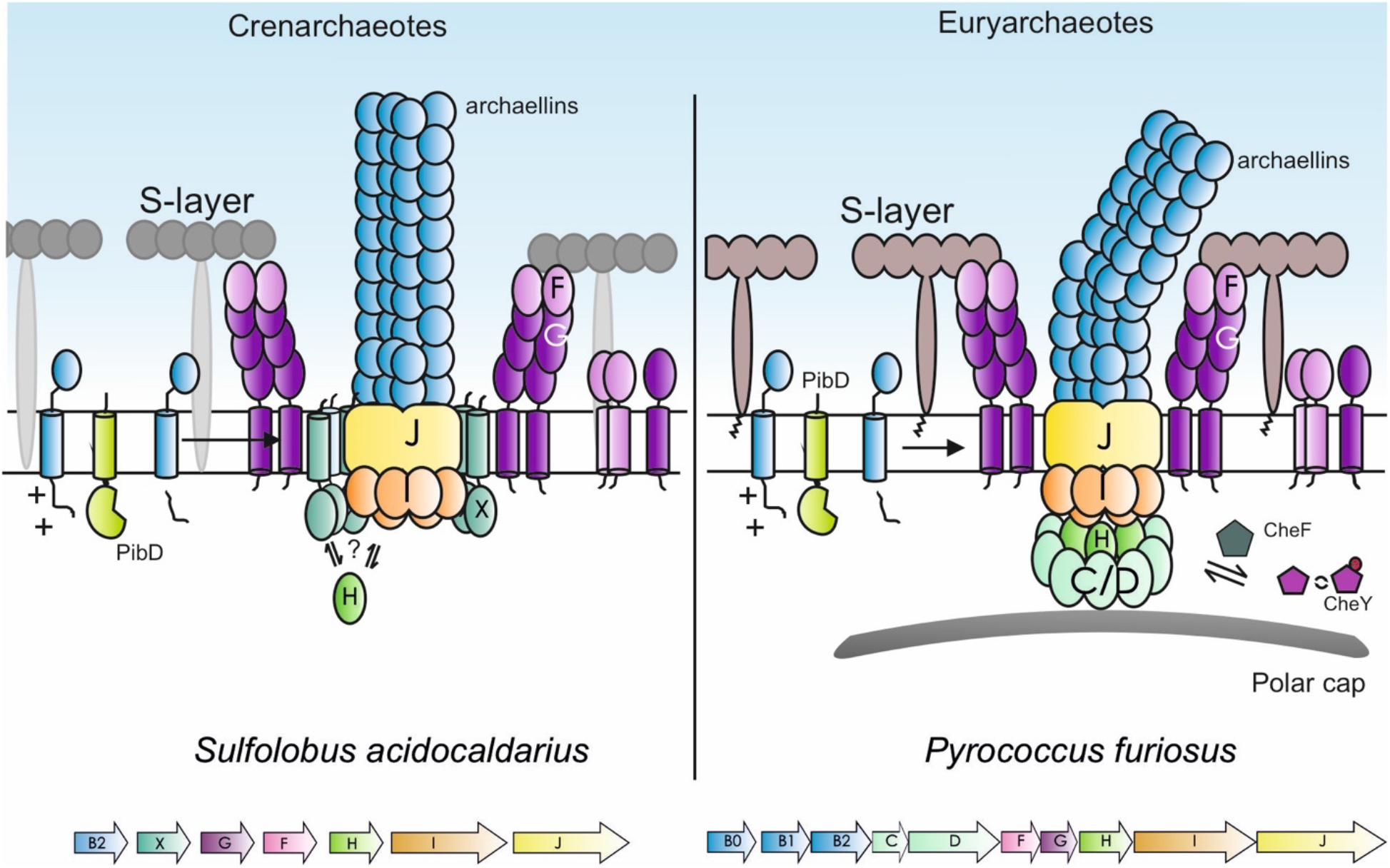
Models of the archaellum assembly system in Cren- and Euryarchaea. Assembly and rotation of archaella in crenarchaeotes is driven by a complex consisting of i) a motor complex formed by the ATPase ArlI, the membrane protein ArlJ and the ATP binding protein ArlH, ii) ArlX which forms a ring around the motor complex and iii) a stator complex formed by ArlF and ArlG. This complex assembles one or more different archaellin proteins into a filament and later also rotates this filament. Archaellins are synthesized as pre-proteins and the membrane embedded prepilin peptidase PibD/ArlK cleaves the signal-peptide prior to the integration of archaellins in the filament. In euryarchaeotes, ArlX is replaced by ArlC/D/E or fusions thereof. ArlC/D/E link the bacterial-like chemotactic machinery to the archaellum motor complex via the archaeal specific adaptor protein CheF and the response regulator CheY. ArlC/D/E have been suggested to interact with the cytosolic polar cap which has only been identified in euryarchaea.

As with any rotary motor, the archaellum motor requires both a rotor and a stator. It has been hypothesised that ArlX, ArlG, and ArlF are part of the crenarchaeal stator (22–24), whereas ArlH, ArlI, and ArlJ form the rotor. ArlG and ArlF form a complex at the tip of a filament formed by ArlG. ArlG is anchored in the membrane and ArlF binds to the S-layer proteins, thus anchoring the archaellum to the S-layer (24). ArlX contains a N-terminal transmembrane domain followed by a cytosolic domain. ArlX from *S. acidocaldarius* (*Sa*ArlX) forms high molecular weight complexes that adopt a ring-like structure large enough to enclose the ArlH/I/J motor complex (22, 25).

ArlI is a hexameric ATPase. Conformational variation in the hexamer of *S. acidocaldarius* ArlI (*Sa*ArlI) provides a model for how successive rounds of ATP binding, hydrolysis and product release result in conformational changes that drive both archaellum assembly and rotation (27). Deletion of the first 29 residues of *Sa*ArlI abolished motility but not archaella assembly, demonstrating that these two processes can be separated (27). The polytopic membrane protein ArlJ, homologous to the platform protein present in T4P (e.g. PilC of *Myxococcus xanthus*), is not only conserved across motile archaea but also essential for archaella assembly (10, 28–31). The structure of ArlJ is unknown, but bioinformatical predictions suggest the presence of positively charged cytosolic loops that may mediate its interaction with ArlI. ArlJ might then transfer the ATP-dependent conformational movements of ArlI across the cell membrane (27, 29). The structures of *Sa*ArlH and ArlH from *Methanocaldococcus jannaschii (Mj*ArlH) have been determined (26, 32). Both proteins crystallised as monomers, but structural modelling based on homologous proteins of known structure suggests that ArlH can also form higher oligomeric species. Indeed, cross-linking experiments showing that *Pf*ArlH has the potential to form hexamers *in vitro* (26, 32). *Sa*ArlH, *Mj*ArlH and *Pf*ArlH bind ATP but none of them showed ATPase activity in standard ATPase assays (26, 32). ArlH possesses the canonical Walker A (WA) as well as a Walker B (WB) motif which is typical of RecA family proteins, where a threonine or serine residue follows the conserved aspartate instead of the carboxylate seen in AAA+ ATPases (26, 32–34).

*Sa*ArlH, *Sa*ArlI and *Sa*ArlX have been shown to interact with each other with high affinity (25) and *Sa*ArlH was imaged by cryo-EM showing 8-10 monomers inside the *Sa*ArlX ring (26). A composite model of the archaellum machinery of *P. furiosus* shows a bell-shaped cytosolic complex of ArlI and ArlH and a surrounding cytosolic ring, most likely consisting of ArlC/D/E (20). Several studies have addressed the interaction between ArlH and ArlI. Mutations in the Walker A and B motifs of *Sa*ArlH are not only critical for ATP binding but also abolish the interaction between *Sa*ArlI and *Sa*ArlH. Similar mutations in *Pf*ArlH abolish the observed stimulation of the ATPase activity of *Pf*ArlI by *Pf*ArlH. Accordingly, *S. acidocaldarius* cells carrying genomic *arlH* WA and WB mutations were unable to assemble archaella, indicating a key regulatory function in archaellum assembly and motility for ArlH (26). Furthermore, purified *Sa*ArlH and *Pf*ArlH retain significant amounts of bound nucleotide and removal of the nucleotide by ammonium sulphate precipitation or introduction of mutations that negatively interfere with ATP binding destabilize these proteins. These data together demonstrate the importance of nucleotide binding to ArlH and its interaction with ArlI. Of note, the interaction between *Mj*ArlH and *Mj*ArlI became weaker with increasing ATP concentration (32).

Both crystal structures of ArlH showed a typical RecA/RAD51 fold with a large, central β-sheet surrounded by α-helices. Interestingly, the closest structural homologue of ArlH is KaiC, the central protein in the regulation of the cyanobacterial circadian rhythm (26, 32, 35). KaiC has two domains (KaiCI and KaiCII) which assemble into a hexameric dumbbell-shaped complex (36). The N-terminal domain, formed by KaiCI, has been associated with hexamerisation of the complex (37), whereas the C-terminus formed by the KaiCII domain shows sequential autophosphorylation and autodephosphorylation at S431 and T432. This sequential phosphorylation and dephosphorylation regulates the periodicity of the cyanobacterial circadian cycle (35, 36, 38–43).

Given the homology between KaiC and ArlH, we set out to further explore the biochemical properties of ArlH and compare them with KaiC. In this study, we show that ArlH is structurally homologous to the KaiCII domain of KaiC and that ArlH undergoes autophosphorylation. In addition, we show that ArlH hexamerises in the presence of ArlI and that this interaction depends on the phosphorylation status of ArlH. Altogether, this allows us to propose a mechanism on how ArlH modulates ArlI via a phosphorylation-dependent mechanism.

## Results

### ArlH is a KaiC-like protein with autophosphorylation activity

Previous studies have shown that ArlH is homologous to KaiC, the central component of the circadian rhythm clock of cyanobacteria (26, 32, 39). Since KaiC has two ATPase domains (KaiCI, involved in hexamerisation and KaiCII, involved on autophosphorylation (39–41)), while ArlH consists of a single ATP-binding domain, the structure of *Sa*ArlH was aligned with the *Synechococcus elongatus* KaiCI and KaiCII domains, as defined by Pattanayek et al., 2004 (36). This structural alignment showed a higher similarity of *Sa*ArlH with KaiCII (rmsd 2.402 Å, 754 atoms) than with KaiCI (rmsd 4.195 Å, 1029 atoms) (Fig. 2). A similar observation was made when *Mj*ArlH was aligned to the KaiCI and KaiCII domains (Supplementary Fig. 1). Since ArlH shows more similarity to the KaiCII domain, this suggested that ArlH might have autophosphorylation activity.

**Figure 2.**
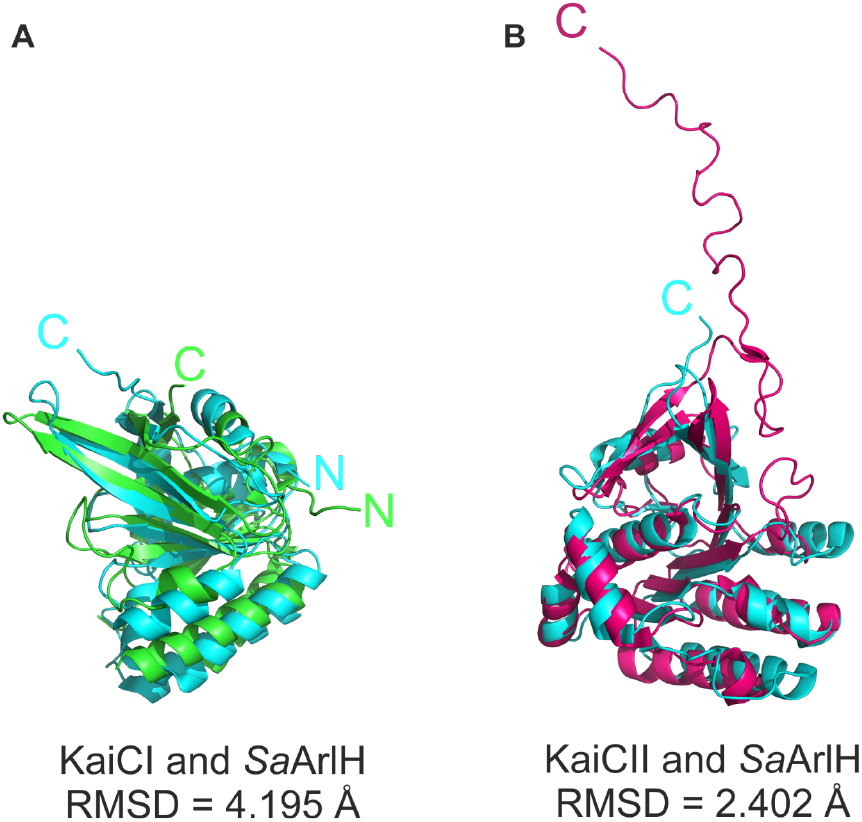
Sa*ArlH is structurally more similar to the KaiCII domain than to KaiCI domain of KaiC*. Structural alignment of (A) *Sa*ArlH (in cyan; PDB: 4YDS) and KaiCI (in green; residues 16-246; PDB: 4TL8) or (B) *Sa*ArlH and KaiCII (in pink; residues 262-519; PDB: 2GBL). For both alignments, solvent molecules, ions, and other ligands were removed prior to structural alignment. The alignment was performed on PyMol with the command “align”, and outlier atoms were removed. identified in euryarchaea.

The hyperthermophile *S. acidocaldarius* has been an important model organism for studies on archaeal motility. When heterologously expressed *Sa*ArlH was purified and incubated with γ-^32^P-ATP in the presence of Mg^2+^ at 55°C for 1 hour, a time-dependent increase in signal was observed at the height of *Sa*ArlH after separation on acrylamide gels (Fig. 3 and Supplementary Fig. 2). No signal was observed when similar assays were performed with *Sa*ArlH and α-^32^P-ATP, demonstrating that the signal is derived from the transferred γ-phosphate. To exclude that the observed signal is derived from a hypothetical contaminating *E. coli* promiscuous protein kinase, a similar phosphorylation assay was performed with *Sa*ArlH variants which were mutated in the Walker A (*Sa*ArlH^K33A^) or the Walker B (SaArlH^D122N^) motifs (Supplementary Fig. 2). Both mutations result in strongly reduced ATP binding (26) and no phosphorylation signal was observed (Fig. 3). Together these data show that *Sa*ArlH undergoes slow time-dependent autophosphorylation *in vitro*. In order to test whether autophosphorylation is a universal property of ArlH proteins, it was tested whether an ArlH from an Euryarchaeal hyperthermophile can also autophosphorylate. Wild-type *Pf*ArlH and *Pf*ArlH with mutations in the WA (*Pf*ArlH^K39A^) and WB (*Pf*ArlH^D126A^) motifs were also purified (Supplementary Fig. 2) and tested in autophosphorylation assays. Similar results to those obtained for *Sa*ArlH were obtained for *Pf*ArlH (Fig. 3), suggesting that autophosphorylation activity is a conserved feature of ArlH.

**Figure 3.**
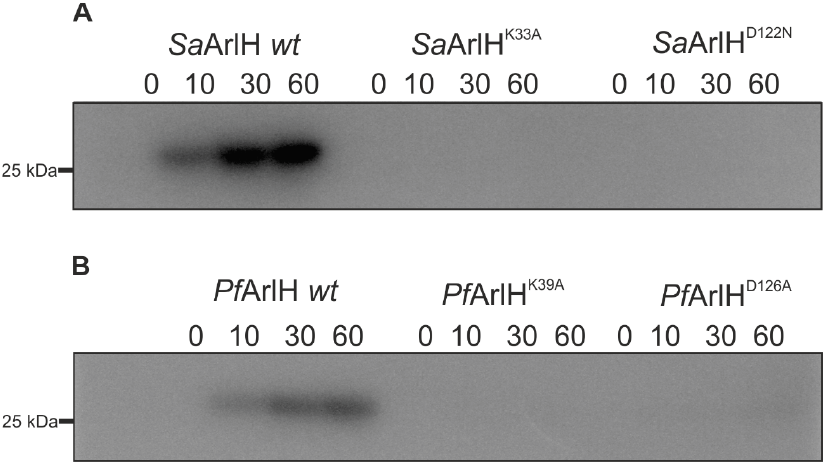
ArlH shows autophosphorylation activity. (A) WT *Sa*ArlH and SaArlH proteins containing mutations in the walker A (K33A) and walker B (D122N) motifs were incubated with γ-^32^P-ATP for different times at 55°C, separated on SDS-PAGE and analysed by phospho-imaging. (B) Similar experiments were performed at 65°C with WT *Pf*ArlH and *Pf*ArlH proteins containing mutations in the walker A (K39A) and walker B (D126A) motifs. The molecular size marker is shown.

### ArlH oligomerises in the presence of ArlI

Although a model of a hexameric ArlH has been proposed, experimental evidence for the oligomeric state of this protein in the presence of other archaellum motor proteins is lacking (26, 32). To study the oligomerization of ArlH in the presence of ArlI, single molecule total internal reflection fluorescence (smTIRF) microscopy was used to explore different conditions under which ArlH might form oligomers. Similar studies have been performed to count the subunits in other systems like, e.g., FliM from flagella and pRNA from the phi29 DNA-packaging motor (44–48). The experiments were performed with single cysteine mutants of *Pf*ArlH (*Pf*ArlH^S109C^) and *Pf*ArlI since these proteins were more stable and higher labelling efficiencies could be obtained than with different *Sa*ArlH and *Sa*ArlI cysteine mutants. The size of the ArlH oligomers was determined by two colour single molecule fluorescence bleaching step assays in the presence of *Pf*ArlI. First, biotinylated *Pf*ArlI was immobilized in a microfluidic chamber (Fig. 4*A*); then, fluorescently labelled *Pf*ArlH was fed into the microfluidic chamber, resulting in immobilisation of *Pf*ArlH via *Pf*ArlI on the surface. Since less bleaching steps (e.g., three instead of six) are easier to detect in a single fluorescence bleaching trace, labelling of *Pf*ArlH was performed with two different dyes, Atto550 and Atto647N. The chamber was alternatingly illuminated with a green (for Atto550) and a red (for Atto647N) laser light and the bleaching steps in each colour were determined (Fig. 4*B*). The summed-up bleaching steps per oligomer were then corrected for the degree of labelling (see methods for details). The results from six different repetitions on six different days with six different sample batches are shown in Fig. 5A for *Pf*ArlH (yellow). Several different oligomer sizes can be observed, including a significant number of hexamers (about 12%) and trimers (about 37%). This shows that ArlH oligomerises in the presence of ArlI.

**Figure 4.**
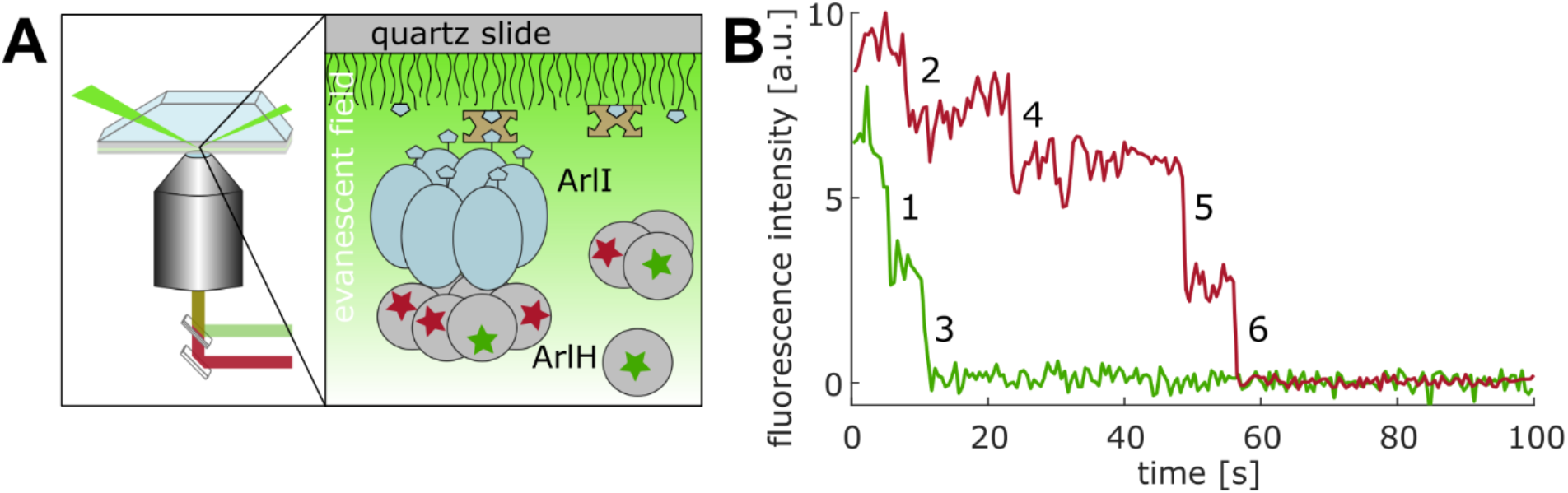
Single-molecule TIRF measurements were performed to assess the oligomeric state of ArlH. A) Schematics of the two colour single molecule TIRF experiment. Biotinylated ArlI is immobilized on the PEGylated quartz slide via biotin-NeutrAvidin binding. Oligomers of labelled ArlH bind to ArlI. B) Example trace with a total of six bleaching steps.

**Figure 5.**
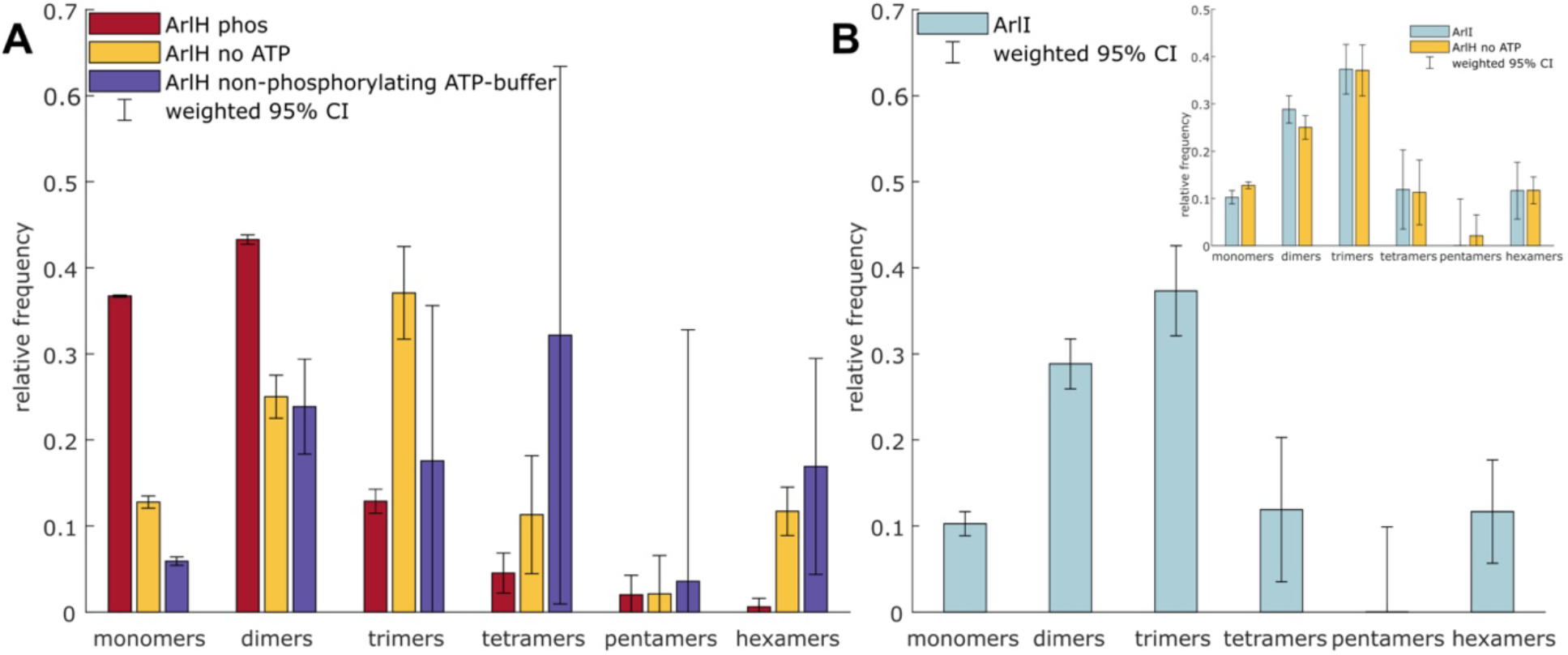
ArlH interacts with ArlI as a hexamer and shifts to lower oligomeric forms when phosphorylated. A) Comparison of the measured oligomer size for *Pf*ArlH without ATP (yellow), phosphorylated *Pf*ArlH (red) and with ATP under non-phosphorylating conditions (violet), showing that ATP binding is not sufficient to regulate *Pf*ArlH’s oligomerization. B) TIRF bleaching experiments with labelled, biotinylated *Pf*ArlI (20 pM) confirm that *Pf*ArlI forms oligomers. In the inset, the bleaching experiments performed on *Pf*ArlI are directly compared with the bleaching experiments performed for *Pf*ArlH without ATP, revealing that the distributions are very similar.

### ArlH autophosphorylation reduces the oligomer size of ArlH in presence of ArlI

To test the effect of ATP on the oligomerization of *Pf*ArlH, the smTIRF bleaching experiment was repeated in the presence of ATP under two different conditions. First, to test for the effect of ATP binding, i.e., in the non-phosphorylating conditions, *Pf*ArlH was incubated with ATP in the presence of Mg^2+^ at room temperature (RT) for 1 hour. Since *Pf*ArlH is derived from a thermophilic organism, at RT, *Pf*ArlH binds ATP with high affinity (Supplementary Fig. 5) but the phosphorylation activity is very low. Second, to test for the effect of ATP hydrolysis, i.e., under phosphorylating conditions, *Pf*ArlH was incubated with ATP in presence of Mg^2+^ at 65°C for one hour. Under non-phosphorylating conditions a clear shift to higher oligomeric species was observed when compared to the absence of ATP, with the two populations differing significantly with a shift to higher oligomeric species under non-phosphorylating conditions (t-test (t(6242)=14.6559, p<<0.001, 6242 = degrees of freedom = sum of oligomers of the two conditions compared here) (Fig. 5A, violet).

In the second case, when ArlH was phosphorylated, the oligomer size clearly decreased to mainly monomers and dimers (altogether 80%) (Fig.5A red). This result is confirmed by a t-test (t(5576)=-29.3370, p<<0.001, 5576 = degrees of freedom = sum of oligomers of the two conditions compared here). Thus, phosphorylated ArlH forms the smallest oligomeric species (mainly monomers and dimers) in the presence of *Pf*ArlI, while non-phosphorylated ArlH already shows larger oligomeric species (more than 50% oligomers larger than dimers), which is further increased if in the presence of excess ATP (but with non-phosphorylated protein).

### *Pf*ArlH interacts with *Pf*ArlI in a 1:1 stoichiometry

Previous data has provided overwhelming evidence that ArlI forms hexamers (20, 27, 32, 49). Moreover, the homologous ATPases PilT and PilB of T4P systems have also been shown to form hexamers (50). Fig. 5B shows the result of two independent TIRF bleaching experiments with labelled *Pf*ArlI^C290S/T411C^. Hexamers can clearly be observed, but also many trimers and dimers. In fact, the oligomer size distribution looks remarkably like that of *Pf*ArlH interacting with *Pf*ArlI (Fig. 5B, inset). The populations of unphosphorylated *Pf*ArlH and *Pf*ArlI show no significant difference (t-test: p=0.7141 t-value (4092)=0.3663). This strongly hints towards a 1:1 binding stoichiometry between *Pf*ArlH and *Pf*ArlI. Therefore, our smTIRF data suggests that ArlH exists as a hexamer at least in one, most likely the functional, form when it interacts with ArlI.

### Phosphorylation inhibits the interaction between *Sa*ArlI and *Sa*ArlH

To test whether phosphorylation also modifies the *Sa*ArlH/*Sa*ArlI interaction, *Sa*ArlH was incubated for 3 hours at 55°C with ATP to allow for autophosphorylation. Then the interaction between ArlI and ArlH was tested using bio-layer interferometry. While non-phosphorylated *Sa*ArlH binds *Sa*ArlI with a K_D_ of 638 ± 161 nM, their interaction is severely reduced when *Sa*ArlH is incubated under phosphorylating conditions prior to the binding assay (Fig. 6). This strongly supports that the phosphorylation status of ArlH universally modulates its interaction with ArlI.

**Figure 6.**
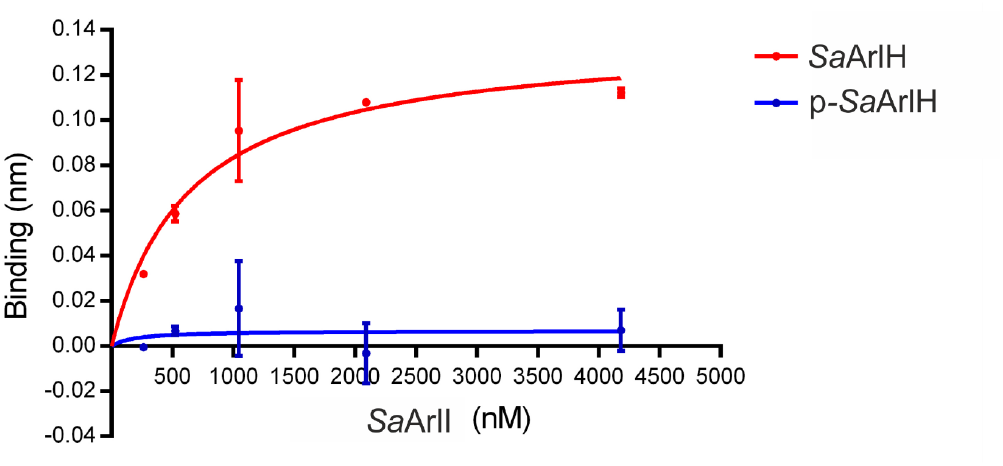
The interaction between SaArlH and SaArlI is strongly reduced when SaArlH is phosphorylated. Bio-layer interferometry assays with *Sa*ArlH and *Sa*ArlI were performed. His-tagged *Sa*ArlH or His-tagged *Sa*ArlH pre-incubated with ATP at 55°C (p-*Sa*ArlH) were bound to the surface and incubated with different amounts of *Sa*ArlI. *Sa*ArlI interacted with surface bound *Sa*ArlH with a K_D_ of 638 ± 16 nM. When *Sa*ArlH was pre-incubated with ATP, thus resulting in phosphorylated *Sa*ArlH, *Sa*ArlI bound *Sa*ArlH with a much lower affinity.

### Identification of phosphorylation sites in ArlH

Since phosphorylation of ArlH might be an important mechanism in the function of the archaellum, identifying the phosphorylated residues was particularly important. Unfortunately, despite extensive efforts, it was not possible to determine the putative phosphorylation sites of ArlH using mass spectrometry. Therefore, different residues which could be involved in phosphorylation were mutated and the resulting protein variant characterised. The *Sa*ArlH T171 and S172 residues, which are located in the same region of the alignment as *S. elongatus* KaiC S431 and T432 (Supplementary Fig. 3), were mutated to alanine. Autophosphorylation assays with the double mutant *Sa*ArlH^T171A/S172A^ showed that phosphorylation was identical to that of the wild-type *Sa*ArlH (Supplementary Fig. 4), demonstrating that these residues do not play a role as phosphorylation sites. Structural alignment of ArlH with the KaiCII domain suggested another Ser/Thr pair (T55 and T56 in *Sa*ArlH) located in the vicinity of the ATP-binding pocket of ArlH. This pair is highly conserved across ArlH proteins, although not in KaiC. These residues were mutated to the structurally related valine amino acid and the purified proteins (Supplementary Fig. 2) analysed for autophosphorylation. While the mutant *Sa*ArlH^T55V^ shows the same phosphorylation as wild-type *Sa*ArlH, phosphorylation was abolished in *Sa*ArlH^T56V^ (Supplementary Fig. 4). A MANT-ATP binding assay was performed with both mutants to check whether they were still able to bind ATP. While *Sa*ArlH^T55V^ still bound ATP, *Sa*ArlH^T56V^ was unable to bind ATP (Supplementary Fig. 5). This explains the absence of phosphorylation in *Sa*ArlH^T56V^, but neither supports nor refutes T56 as a potential phosphorylation site. Similar residues in *Pf*ArlH (S61 and S62) were mutated to the structurally similar cysteine residue, and these variants were purified (Supplementary Fig. 2). In these cases, all three variants -wild-type and single mutants for each residue -showed similar autophosphorylation patterns (Supplementary Fig. 4) and similar ATP-binding affinities (Supplementary Fig. 6), suggesting that at least in *Pf*ArlH these residues are not involved in phosphorylation.

### The conserved catalytic glutamate is important for ArlH autophosphorylation and essential for motility

Since no further clear candidates for the phosphorylation site in ArlH could be identified by bioinformatics, the conserved E57 was mutated. This residue is conserved across ArlH proteins, except for several Euryarchaeota, where the residue at this position is a glutamine instead of a glutamate. This residue aligns with two conserved glutamate residues (E318 and E319 in *S. elongatus*) which have been proposed to play an important role in phosphorylation, with E318 functioning as the general base (33, 36–38, 41, 51). Mutations in this residue in KaiC either abolish or reduce its autophosphorylation activity (37, 38, 52). This residue was mutated in both *Sa*ArlH and *Pf*ArlH. *Sa*ArlH^E57A^, *Sa*ArlH^E57Q^ (Fig. 7*A*) and *Pf*ArlH^Q63E^ (Fig. 7B) showed lower autophosphorylation activity than the wild-type protein, whereas *Pf*ArlH^Q63A^ showed wild-type levels of phosphorylation (Fig. 7*B*). It is difficult to test motility of *P. furiosus* as it is a hyperthermophilic anaerobic organism, but genomic *SaarlH*^*E57A*^ and *SaarlH*^*E57Q*^ single mutants were created and tested for both archaellation and motility on semi-gelrite plates at 75°C (Fig. 8). Strikingly, neither mutant is archaellated nor motile (Fig. 8), suggesting that optimum phosphorylation is essential for archaellation.

**Figure 7.**
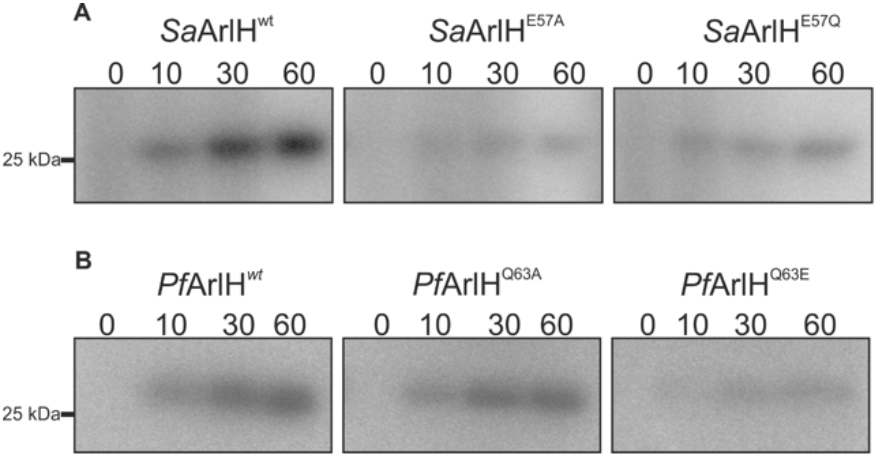
The autophosphorylation activity of ArlH is reduced when the catalytic residue is mutated to an amino acid of opposite charge. The catalytic glutamate of KaiCII is conserved in ArlH, with the exception of Euryarchaeota which seem to have instead an invariable glutamine at this position. (A) WT *Sa*ArlH and *Sa*ArlH proteins containing mutations in the catalytic glutamate (E57A and E57Q) were incubated with γ-^32^P-ATP for different times at 55°C, separated on SDS-PAGE and analysed by phospho-imaging. (B) Similar experiments were performed at 65°C with WT *Pf*ArlH and *Pf*ArlH proteins containing mutations in the corresponding glutamine (Q63A and Q63E).

**Figure 8.**
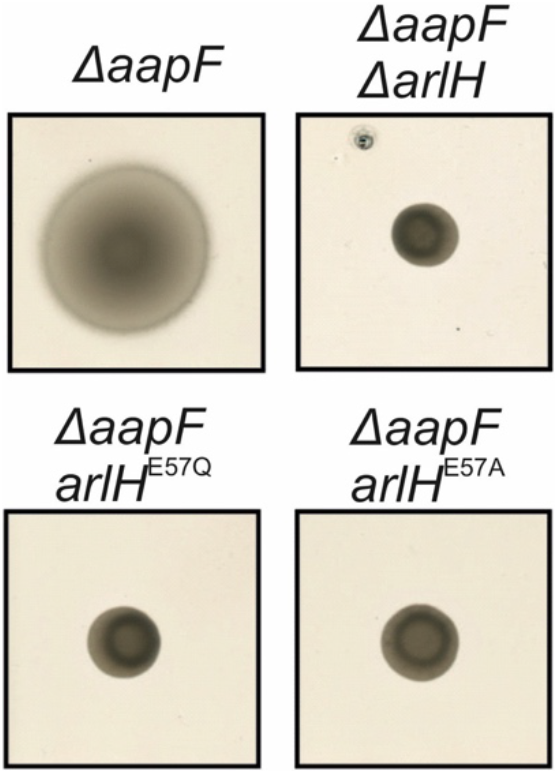
Optimal phosphorylation levels are essential for motility. The motility of *S. acidocaldarius* strains carrying the indicated genomic mutations was determined. Same amounts of cells were spotted on semi-solid gelrite plates. After incubating at 75°C for 5 days, the plates were scanned and recorded. Δ*aapF* and Δ*aapF*Δ*arlH* were used as hyper-motile and non-motile control, respectively. The figures shown here were representative examples of biological triplicates.

## Discussion

KaiC-like proteins have been identified in Archaea, where they were recognised as part of the superfamily that includes DnaB from bacteria, Rad51/DMC1 from eukaryotes, and RadA from archaea (33, 53). KaiC proteins were first identified in cyanobacteria (54). In these organisms, the rhythmic phosphorylation and dephosphorylation of KaiC regulates global gene expression in a circadian manner, allowing the cells to anticipate periods of light and dark and adapt their metabolism, e.g., photosynthesis and nitrogen fixation, accordingly (35). KaiC-like proteins in other organisms have been assumed to function in similar processes. However, bioinformatic analyses have proposed that KaiC-like proteins in archaea underwent several instances of functionalisation and might be involved in signal transduction (53, 55). Indeed, there already is evidence for the functional versatility of KaiC proteins. For example, a KaiC homologue from *Legionella pneumophila* contributes to oxidative and ionic stress resistance (56), but does not function as a clock and does not interact with an also present KaiB homologue (56). In archaea, KaiC-like proteins have been proposed to interact with putative output domains, of which the ArlH-ArlI interaction is an example (55).

ArlH is a single-domain KaiC-like protein. A previous analysis based on sequence alignment suggested archaeal KaiC-like proteins to be more similar to the KaiCI domain (57). Here, however, structural alignments with *Mj*ArlH and *Sa*ArlH structures reveal a higher similarity with KaiCII. Indeed, autophosphorylation takes place in the KaiCII domain and this and previous studies (56, 58, 59) have shown that KaiC-like proteins can autophosphorylate, suggesting that autophosphorylation is a property of KaiC-like proteins independent of their specific function. In this study, a relatively low level of phosphorylation was observed in the autoradiography assays. This might be caused by i) some ArlH being purified in a nucleotide-bound state and a slow release rate for the nucleotide ii) lack of a component that stimulates the phosphorylation rate or iii) ArlH being both an auto-kinase and an autophosphatase, with these antagonistic activities resulting in a dynamic equilibrium between phosphorylated and non-phosphorylated ArlH.

Previous studies on cyanobacterial KaiC and members of the RecA family demonstrated that ATP hydrolysis by proteins with a RecA-like fold requires residues from adjacent RecA-like domains and that hexamerisation is a common theme in this group of proteins (60, 61). This was also suggested for the KaiC-like proteins PH0186 and PH0284 from *Pyrococcus horikoshii* OT3, while another KaiC-like protein from the same organism forms tetramers (62– 64). ArlH was suggested to fulfil its function in the archaellum motor as a hexamer and a model based on this hypothesis was proposed (26). Previous evidence for different oligomeric forms of ArlH had been obtained (26) and the data presented in this study further confirms and proposes that ArlH interacts with ArlI as a hexamer. Using smTIRF bleaching experiments, *Pf*ArH is shown to exist as higher oligomeric species in the presence of *Pf*ArlI. The frequency distribution of bleaching steps for *Pf*ArlH is very similar to the frequency distribution of bleaching steps for *Pf*ArlI, a protein that has been extensively shown to form hexamers (27, 49, 65). This and data suggests that *Pf*ArlH and *Pf*ArlI interact in a 1:1 stoichiometry and, therefore, that *Pf*ArlH forms hexamers (65).

ArlH has previously been show to form monomers in solution, although cross-linking experiments suggest as well that ArlH has the potential to form hexamers (26, 32). The ArlI requirement for stable oligomerisation bears similarities with other RecA proteins which have been reported to require additional domains for a stable hexamerisation (61). Hexamerisation of KaiC is mainly driven by the KaiCI domain, with the KaiCII having only a stabilising effect on the hexamer (38). The observation of lower-order oligomers in these experiments is likely due to the low protein concentrations (pM range) to which the proteins are diluted shortly before the single-molecule experiments. These concentrations might be around or slightly below the K_D_ for the monomers’ interaction for both ArlH and ArlI.

Once the ArlI-dependent oligomerisation of *Pf*ArlH had been established, the impact of *Sa*ArlH phosphorylation on the *Sa*ArlH-*Sa*ArlI interaction was assessed. Biolayer interferometry measurements showed that when *Sa*ArlH is phosphorylated, the interaction between *Sa*ArlI and *Sa*ArlH is strongly reduced (Fig. 6). smTIRF experiments in the presence of *Pf*ArlI revealed that when *Pf*ArlH is phosphorylated there is a shift to lower oligomeric species when compared to non-phosphorylated *Pf*ArlH. Such a shift was not observed when assays were performed in the presence of ATP under non-phosphorylating conditions, demonstrating that phosphorylation –not nucleotide binding – is responsible for this change in the oligomeric state. Phosphorylation thus seems to be a conserved feature that modulates the interaction between ArlH and ArlI.

Several attempts at characterising which residues are phosphorylated in this protein failed, as no phospho-peptides could be detected by mass-spectrometry for either *Sa*ArlH or *Pf*ArlH. It was however possible to identify a residue – E57 in *S. acidocaldarius* – which results in reduced autophosphorylation activity when mutated to either an alanine or a glutamine residue. Intriguingly, the homologous residue in *P. furiosus* is a glutamine instead of a glutamate – a trend that seems to be common in Euryarchaea apart from halophiles. Mutating the glutamine residue to an alanine does not interfere with the autophosphorylation of *Pf*ArlH but changing this residue to a glutamate results in lower phosphorylation activity. Insertion of these mutations in *S. acidocaldarius* cells completely abolished archaella synthesis and, consequently, motility. These results strongly imply that optimum phosphorylation activity of ArlH is a strict requirement for archaella assembly/maintenance. Moreover, the data suggests that despite their presumably similar functions there are mechanistic differences between ArlHs of cren-and euryarchaeotes.

Together with the results from previous studies (26, 65) we can now describe the interaction between ArlH and ArlI in more detail, namely: i) ATP binding to ArlH is a requirement for the interaction with ArlI; ii) ArlH interacts with ArlI as a hexamer; iii) hexamerisation of ArlH is ArlI-dependent and iv) phosphorylation of ArlH modulates the ArlH-ArlI interaction. Thus, both the nucleotide dependent ArlH-ArlI interaction and modulation of this interaction by phosphorylation of ArlH is likely required in early stages of archaella assembly. This is further supported by the absence of archaella filaments when cells express ArlH mutated in the Walker motifs or with mutations in the E57 residue, which reduce autophosphorylation. To understand how the interaction between ArlH and ArlI and its modulation by phosphorylation of ArlH exactly drives the assembly and, at later stages, the rotation of the archaellum must be studied in the presence of other components of the archaellar motor like ArlJ and ArlX (or ArlC/D/E), as these proteins interact with and/or modulate ArlH and ArlI.

## Experimental procedures

### Strains and Plasmids

Strains, plasmids and primers used in this study can be found in **Tables 1, 2** and **3** respectively, in the Supporting Information. *E. coli* TOP10 (Invitrogen) was used for cloning and plasmid amplification and *E. coli* Rosetta cells were used for heterologous expression. The NEB ER1821 *E. coli* strain was used to methylate plasmids for transformation into *S. acidocaldarius*. Heterologous overexpression of proteins in *E*.*coli* took place from pETDuet1 derived plasmids with the gene of interest cloned in the multiple cloning site. Insertion of tags was performed by *in vivo* DNA assembly (66). Site-directed mutagenesis was performed by round PCR using the indicated primers. Insertion of the AviTag (GLNDIFEAQKIEWHE) upstream of *arlI* in pSVA3116 was performed via an intermediate plasmid containing half of the tag, followed by insertion of the remaining fragment using the two indicated primer pairs. All plasmid sequences were confirmed by sequencing.

*In vivo* point mutations in *S. acidocaldarius* were generated following the protocol described by Lassak *et. al* (10) with a modification. Shortly, the *arlH* gene was amplified by PCR from the genomic DNA of *S. acidocaldarius* DSM639, including 700 bp up-and downstream regions. The primers used for amplification were 3614/3617. The fragment was cloned as an ApaI/PstI digested fragment into the pSVA406 vector, yielding plasmid pSVA2126. Site directed mutagenesis was performed by round PCR using the plasmid pSVA2126 as template, with the primers indicated in **Table 3** and yielding the plasmids in **Table 2** (Supporting information). The plasmids were methylated in *E. coli* strain ER1821 and transformed into the MW455 *S. acidocaldarius* strain. This strain is a deletion mutant of *arlH* and insertion of the mutated plasmids resulted in knock-in of the *arlH* variants by homologous recombination to its original locus in the genome.

### Cell growth

*Sulfolobus acidocaldarius* DSM639 was grown aerobically at 75°C in Brock basal medium (67) supplemented with 0.1% (w/v) tryptone (Roth) or NZ-Amine AS (Sigma) and 0.2% (w/v) dextrin with pH adjusted to 3.5 with sulfuric acid. All the uracil auxotrophic stains of ArlH point mutations, hyperarchaellated wildtype MW156 (*S. acidocaldarius* Δ*aapF*), non-archaellated MW455 (*ΔarlH* in *S. acidocaldarius* Δ*aapF* background) and the point mutant strains (MW480 and MW490 for the mutations *arlH*^E57Q^ and *arlH*^E57A^, respectively, in the *S. acidocaldarius* Δ*aapF* background) (10) were grown in Brock medium supplemented with 10 μg/ml uracil. To prepare solid medium for *S. acidocaldarius*, Brock medium was solidified with a final concentration of 0.6% (w/v) Gelrite (Sigma), 10 mM MgCl_2_ and 3 mM CaCl_2_ and the pH was adjusted accordingly.

For overexpression in *E*.*coli*, overexpression plasmids were transformed into *E. coli* Rosetta (DE3) cells, which were grown for ∼ 16h at 37°C in LB medium containing ampicillin (100 µg/ml) and chloramphenicol (30 µg/ml). Four litres of LB medium supplemented with ampicillin and chloramphenicol (100 µg/ml and 30 µg/ml, respectively) were inoculated with the pre-culture and grown at 37°C until an OD_600_ of 0.6-0.8. The cultures were induced with isopropyl β-D-thiogalactopyranoside to a final concentration of 0.3 mM. Cells overexpressing biotinylated *Pf*ArlI were further supplemented with D-biotin to a final concentration of 50 µM. Growth was continued for 4h at 37°C for *Pf*ArlH, or for 16h at 18°C for *Sa*ArlH, Strep-*Sa*ArlI, and biotinylated *Pf*ArlI. Cells were harvested by centrifugation and the pellets were either used immediately or stored at −20°C. Cell pellets derived from cultures overexpressing S*a*ArlH and *Pf*ArlH were suspended in 35 ml of buffer A (50 mM HEPES pH 7.0, 150 mM NaCl) while cultures overexpressing Strep-*Sa*ArlI and biotinylated *Pf*ArlI were suspended in buffer B (50 mM Tris pH 8.0, 150 mM NaCl) plus 0.5% Triton X-100 and one tablet of cOmplete ™ Protease Inhibitor Cocktail Tablet (Roche).

### Protein purification

When frozen, the pellets were thawed on ice. The suspensions in either Buffer A or Buffer B (for ArlH and ArlI, respectively) were vortexed until homogeneity, a small amount of DNAse I was added and the cells were disrupted three times in a LM10 Microfluidizer® device at 1500 bar. Cell debris was removed by centrifugation at 4500 g for 20 minutes. The supernatant was further centrifuged at 27200 g for 30 minutes. HIS-Select® Nickel Affinity resin (Sigma) and Streptactin column material (IBA GmbH, Göttingen, Germany) were equilibrated with buffer A and B respectively. The clarified lysate of cells overexpressing *Sa*ArlH, *Pf*ArlH or biotinylated *Pf*ArlI were subsequently mixed with HIS-Select® Nickel Affinity resin (SIGMA) while lysate of cells overexpressing Strep-*Sa*ArlI was mixed with Streptactin column material (IBA GmbH, Göttingen, Germany) and incubated at 4°C with rotation for 1 hour, after which the mixture was poured into a gravity column. The flow-through was collected and the column was washed with 10 column volumes (CV) of 50 mM HEPES pH 7.0, 500 mM NaCl, 20 Mm imidazole and 20 CV of 50 mM HEPES pH 7.0, 150 mM NaCl, 20 mM imidazole for the HIS-Select® Nickel Affinity resin and with 30 CV of 50 mM Tris pH 8.0 for the Streptactin column material. Proteins were eluted in 6 fractions of 1 ml. *Sa*ArlH was eluted in 50 mM MES pH 6.0, 150 mM NaCl, 200 mM imidazole; *Pf*ArlH was eluted in 50 mM HEPES pH 7.0, 150 mM NaCl, 200 mM imidazole; Strep-*Sa*ArlI was eluted in 50 mM HEPES pH 8.0, 150 mM NaCl, 2.5 mM desthiobiotin; and biotinylated *Pf*ArlI was eluted in 50 mM HEPES pH 8.0, 150 mM NaCl, 200 mM imidazole. Imidazole was removed from purified *Sa*ArlH and *Pf*ArlH by overnight dialysis against their respective elution buffers without the imidazole and *Pf*ArlI and biotinylated *Pf*ArlI were further purified by size exclusion chromatography using a Superdex 200 Increase 10/300 GL column equilibrated with 50 mM HEPES pH 8.0, 150 mM NaCl.

### *In vitro* phosphorylation assays

ArlH was added to a concentration of 5 µM in the presence of 0.01 mCi (at least 17 nM) γ-^32^P-ATP (or α-^32^P-ATP) in a final volume of 100 µl of 50 mM MES pH 6.0, 150 mM NaCl with 5 mM MgCl_2_ for *Sa*ArlH or, for *P. furiosus* proteins, buffer B with 5 mM MgCl_2_. Reaction mixtures were incubated for *S. acidocaldarius* proteins at 55°C and for *P. furiosus* proteins at 65°C. At each time-point, 12 µl were taken from each reaction mixture and mixed with an equal volume of 2x concentrated Laemmli buffer. Samples were separated by SDS-PAGE and the radioactive signal was detected by autoradiography using the Typhoon FLA 9500 system.

### Motility assays

The swimming assays were performed on semi-solid (0.15%) gelrite plates following the protocols described previously (10). Cells were grown until an OD_600_ of 0.4-0.5. Samples of the cultures (5 µl) were spotted on the gelrite plates, which were incubated for 4-5 days in a humid chamber at 75°C.

### Biolayer interferometry

The Blitz® system (Pall Life Sciences) was used to measure the interaction of *Sa*ArlH and *Sa*ArlI. NiNTA biosensors (Pall Life Sciences) were pre-hydrated in the assay buffer (20 mM Tris HCl pH 8.0; 150 mM NaCl; 0.2% Tween-20) for 10 mins. Using the Blitz, biosensors were first equilibrated in the assay buffer for 30 s, followed by incubation with 1 µM *Sa*ArlH in assay buffer for 30 s and equilibration in assay buffer for 600 s to establish a stable baseline. Then, the *Sa*ArlH-bound biosensors were dipped in assay buffer containing *Sa*ArlI for 300 s to allow interaction. New NiNTA biosensors were used for each concentration of *Sa*ArlI (261-4181 nM). Signals were corrected for a-specific binding of *Sa*ArlI to the biosensors in the absence of *Sa*ArlH. Binding at 300 s was plotted against the *Sa*ArlI concentration and fitted with the one site specific binding model (B = B_max_ * [SaArlI] / (K_D_ + [SaArlI]), where B = binding [nm]; B_max_ = binding at 300 s [nm]; K_D_ = equilibrium constant [nM]; [SaArlI] = SaArlI concentration) to determine the K_D_.

### ATP binding assays

Five micromolar ArlH in 150 µl 50 mM MES pH 6.0, 150 mM NaCl with 5 mM MgCl_2_ for *S. acidocaldarius* or buffer B with 5 mM MgCl_2_ for *P. furiosus* proteins was titrated with a MANT-ATP stock solution containing 5 µM ArlH. Fluorescence intensity was recorded at excitation and emission wavelengths of 350 and 440 nm, respectively, with slit widths of 2 nm in a temperature-controlled FluoroMax-4 spectrofluorometer (Horiba) at 20°C. The fluorescent signal of MANT-ATP in the absence of protein was determined and subtracted and the fluorescence increase was plotted.

### ArlH labelling

*Pf*ArlH^S109C^ was labelled with maleimide fluorescent dyes. ArlH was incubated with 10 mM tris(2-carboxyethyl)phosphine in a volume of 200 µl and a concentration of about 120 mM for 20 minutes at room temperature. TCEP was removed by concentrating ArlH in PBS buffer (100 mM, pH 6.7) in centrifugal filters (Amicon® Ultra, cutoff 10K) to 150 µl and then washed, in the same centrifugal filter, for five times. The protein was then incubated with up to two fluorescent dyes (1.5-fold molar excess each) over night at 4°C. Atto550 and Atto647N (ATTO-TEC) were used. Afterwards, free dye was removed using a PD 25 MiniTrapTM (GE Healthcare) equilibrated in buffer A. The degree of labelling (DOL) was measured on a NanoDrop (Thermo ScientificTM). In case the DOL did not exceed 50%, the protein was labelled once again for two hours at room temperature. *Pf*ArlIT411C was labelled in a manner similar to *Pf*ArlH, but in 50 mM HEPES pH 8.0, 150 mM NaCl buffer.

### Single molecule TIRF experiments

The experiments were carried out in a custom-built prism type TIRF setup using two lasers at wavelengths 532 nm (green, Coherent® OBISTM LS) and 637 nm (red, Coherent® OBISTM LX) (68). The excitation beam is focused on a quartz prism and onto the flow chamber, which is passivated with a mixture of methoxy-silane-polyethylene glycol (5000 Dalton, Rapp Polymere) and methoxy-silane-PEG-Biotin (3000 Dalton, Rapp Polymere) in an 80:3 ratio. Incubation with BSA solution for 30 minutes prior to measurement (0.5 mg/mL in HEPES buffer, Carl Roth) further improved the passivation. To allow the biotinylated protein sample to bind to the surface, the chamber was further coated with NeutrAvidin (0.25 mg/mL, Thermo Fisher Scientific) and washed with HEPES buffer.

To test the oligomerisation of ArlH in presence of ArlI, biotinylated ArlI (10 nM) was flushed into the flow chamber and incubated for 10 minutes. Unbound sample was washed with buffer, before labelled ArlH (20 pM) was added. After an incubation time of another 10 minutes, excess protein was washed out with HEPES buffer.

For the autophosphorylation experiments, ArlH (10 µM) was incubated with a hundredfold excess of ATP (1 mM) for one hour at 65°C in HEPES buffer with MgCl_2_ (5 mM) prior to measurement. This procedure was also applied for the ATP-binding experiments, however, at room temperature.

To check the oligomer size of ArlI, the procedure varied slightly: as the biotinylated ArlI was labelled with fluorescent dye, only 20 pM were flushed into the chamber to allow for single-molecule measurements.

The emitted fluorescence of the labelled sample is collected by an oil immersion objective (100x magnification, Nikon), separated according to wavelength and recorded by an EMCCD camera with 2×2 binning (iXon Ultra 897, Andor). The recorded movies are saved in the form of 16 bit TIFF stacks, which store the fluorescence intensities of the respective detection channels as successive frames (68).

The measurements were conducted using alternating laser excitation (ALEX) with an excitation time of 200 ms and a dark time of 50 ms, whereby the two alternating wavelengths depended on the fluorescent dyes attached to the protein.

#### Data analysis of smTIRF experiments

Data selection was done in Igor Pro 6.37, using an in-house script. It identifies the positions of single molecules by selecting the brightest spots in the respective detection channel, summing up five consecutive frames. Prior to our experiments, the green and red channel were aligned by image registration with fluorescent beads (TetraSpeck™ microspheres, 0.2 µm, Invitrogen™). Thus, fluorescence intensity traces in all excitation colors per single molecule could be obtained. From those, single molecule bleaching step traces were selected for analysis when having at least one bleaching step. To be recognised as such, the following three criteria must be met. First, a sudden drop in fluorescence intensity is not followed by another increase (blinking) but instead leads to a plateau. Second, in case of more than one bleaching step per trace, the plateaus have roughly the same height. Third, said plateaus are at least two frames (which equals 0.7s) long. Applying these criteria, the bleaching steps in each trace were determined and counted by eye by two independent researchers. This procedure was facilitated by using the fit generating function of AutoStepfinder (69). For each trace, fits displaying varying numbers of bleaching steps were created by adapting the parameters. By eye, the best fit was chosen. When measuring sample with two different fluorescent labels, one oligomer was characterized by two traces. Thus, the steps of both dyes were summed up to obtain the total number of bleaching steps. Afterwards, a histogram showing the absolute frequency per bleaching step was created, as well as another one showing the relative distribution. To correct for the degree of labelling (DOL), the latter was then fitted using a binomial distribution

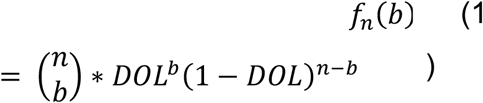

with n being the oligomer size and b the observed bleaching steps. As zero bleaching steps cannot be observed in our experiments, the binomial distributions were truncated:

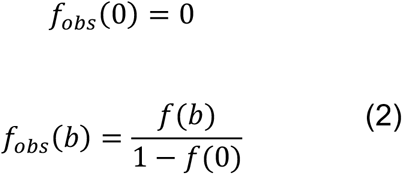

For a mixture of oligomer sizes ranging from monomers to hexamers, the distribution was given by a sum of binomials

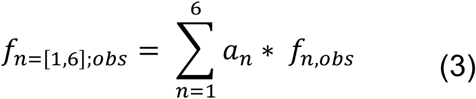

With

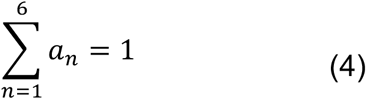

The values of the factors a_n_ are obtained from the fitting procedure and correspond to the relative occurrence of each oligomer.

Every experiment was repeated at least once, however, the number of recorded traces per repetition varied between 173 and 995 traces. To create a joint bar chart in which each oligomer is weighted equally, the absolute frequencies per oligomer size and repetition were cumulated for all experiments. 95% confidence intervals were given by the fitting procedure for each repetition. They were weighted as well according to the number of recorded traces to obtain uncertainties for the joined bar chart. To determine whether the means of two different conditions (e.g. oligomer size of phosphorylated ArlH compared to oligomer size of unphosphorylated ArlH) differ significantly, a two-sample t-test was applied as our datasets are large enough (1906 traces for phosphorylated ArlH, 3671 traces for unphosphorylated ArlH and 2572 traces for ATP-binding, 423 for ArlI) to fulfil the central limit theorem.

## Data availability

All data described in the manuscript are presented in the main text, figures, and supporting information.

## Supporting information

This article contains supporting information.

## Acknowledgments

We thank Tomasz Neiner for initiating the measurements on autophosphorylation of ArlH.

## Author contributions

NM designed the experiments, performed and analysed experiments, and wrote the manuscript. LV and JS performed the experiments and analysed the data. RK and PC performed experiments. CvdD, TH and S.-V.A designed the experiments and wrote the manuscript. All authors read and contributed for the manuscript.

## Funding and additional information

NM, LV, and JS were supported by the Collaborative Research Centre SFB1381 funded by the Deutsche Forschungsgemeinschaft (DFG, German Research Foundation) – Project-ID 403222702 – SFB 1381. NM was supported in part by the Excellence Initiative of the German Research Foundation (GSC-4, Spemann Graduate School) and in part by the Ministry for Science, Research and Arts of the State of Baden-Wuerttemberg.

