## Supporting Information for "Autophosphorylation of the KaiC-like protein ArlH inhibits oligomerisation and interaction with ArlI, the motor ATPase of the archaellum"

**Supplementary tables**

**Table 1 – Strains of *S. acidocaldarius* and *E. coli* used in this study**

| Strains | Relevant characteristics | Source |
|---|---|---|
| ***E. coli*** | | |
| Rosetta (DE3) | $F^-$ *ompT hsdS*$_B$($r_B^-$ $m_B^-$) *gal dcm* (DE3) pRARE ($Cam^R$) | Novagen |
| ER1821 | $F^-$ *glnV44 e14*$^-$*(McrA*$^-$*) rfbD1? relA1? endA1 spoT1? thi-1* Δ*(mcrC-mrr)114::IS10* | NEB |
| TOP10 | $F^-$*mcr*A Δ*(mrr-hsd*RMS-*mcr*BC) φ80*lac*ZΔM15 Δ*lac*X74 *rec*A1 *ara*D139 Δ(*ara-leu*)7697 *gal*U *gal*K *rps*L ($Str^R$) *end*A1 *nup*G | Invitrogen |
| ***S. acidocaldarius*** | | |
| DSM639 | Wild-type *Sulfolobus acidocaldarius* | DSMZ |
| MW001 | DSM 639 Δ*pyrE* | (Wagner et al., 2012) |
| MW156 | MW001 Δ*aapF* (*Saci_2318*) (hypermotile strain) | (Lassak et al., 2012b) |
| MW455 | MW156 Δ*aapF* Δ*arlH* (Saci_2318, *Saci_1174*) (non-motile strain) | (Lassak et al., 2012b) |
| MW480 | MW156 Δ*aapF* Δ*arlH* (Saci_2318, *Saci_1174*)::*arlH*$^{E57Q}$ | This study |
| MW490 | MW156 Δ*aapF* Δ*arlH* (Saci_2318, *Saci_1174*)::*arlH*$^{E57A}$ | This study |

**Table 2 - Plasmids**

| Plasmids | Relevant characteristics | Source |
|---|---|---|
| pSVA2100 | pETDuet-1-based plasmids for the overexpression of $His_6$-*Sa*ArlH expression in *E. coli*. | (Chaudhury et al., 2015) |
| pSVA2130 | pETDuet-1-based plasmids for the overexpression of $His_6$-*Sa*ArlH$^{K33A}$ expression in *E. coli*. | (Chaudhury et al., 2015) |

| | | |
|---|---|---|
| pSVA2131 | pETDuet-1-based plasmids for the overexpression of $His_6$-*Sa*ArlH$^{D122N}$ expression in *E. coli*. | (Chaudhury et al., 2015) |
| pSVA2167 | pETDuet-1-based plasmids for the overexpression of $His_6$-*Pf*ArlH expression in *E. coli*. | (Chaudhury et al., 2015) |
| pSVA2176 | pETDuet-1-based plasmids for the overexpression of $His_6$-*Pf*ArlH$^{K39A}$ expression in *E. coli*. | (Chaudhury et al., 2015) |
| pSVA2178 | pETDuet-1-based plasmids for the overexpression of $His_6$-*Pf*ArlH$^{D126A}$ expression in *E. coli*. | (Chaudhury et al., 2015) |
| pSVA406 | Amp$^r$, gene targeting plasmid, pGEM-T Easy backbone, pyrEF cassette of *S. solfataricus* | (Wagner et al., 2012) |
| pSVA2126 | Amp$^r$, *arlH* cloned into pSVA406 with ApaI-PstI | (Chaudhury et al., 2016) |
| pSVA3157 | Amp$^r$, *arlH* E57Q was created by round pcr on pSVA2126 by using primer pairs 5167 and 5168 | This study |
| pSVA3188 | Amp$^r$, *arlH* E57A was created by round PCR on pSVA2126 by using primer pairs 3653 and 3654 | This study |
| pSVA2156 | pETDuet-1-based plasmids for the overexpression of $His_6$-*Sa*ArlH$^{E57A}$ expression in *E. coli*. | This study |
| pSVA3156 | pETDuet-1-based plasmids for the overexpression of $His_6$-*Sa*ArlH$^{E57Q}$ expression in *E. coli*. | This study |
| pSVA5548 | pETDuet-1-based plasmids for the overexpression of $His_6$-*Pf*ArlH$^{Q63A}$ expression in *E. coli*. | This study |
| pSVA5549 | pETDuet-1-based plasmids for the overexpression of $His_6$-*Pf*ArlH$^{Q63E}$ expression in *E. coli*. | This study |

| | | |
|---|---|---|
| pSVA2160 | pETDuet-1-based plasmids for the overexpression of His$_6$-*Sa*ArlH$^{T171A/S172A}$ in *E. coli*. | This study |
| pSVA5529 | pETDuet-1-based plasmids for the overexpression of His$_6$-*Sa*ArlH$^{T55V}$ expression in *E. coli*. | This study |
| pSVA5530 | pETDuet-1-based plasmids for the overexpression of His$_6$-*Sa*ArlH$^{T56V}$ expression in *E. coli*. | This study |
| pSVA13307 | pETDuet-1-based plasmids for the overexpression of His$_6$-*Pf*ArlH$^{S61C}$ expression in *E. coli*. | This study |
| pSVA13308 | pETDuet-1-based plasmids for the overexpression of His$_6$-*Pf*ArlH$^{S62C}$ expression in *E. coli*. | This study |
| pSVA3116 | pETDuet-1-based plasmid for the overexpression of His$_6$-*Pf*ArlI expression in *E. coli*. | (Chaudhury et al., 2015) |
| pSVA5500 | pETDuet-1-based plasmid for the overexpression of His$_6$-AviTag-*Pf*ArlI expression in *E. coli*. | This study |
| pSVA5811 | pETDuet-1-based plasmid for the overexpression of His$_6$-*Pf*ArlH$^{S109C}$ expression in *E. coli*. | This study |
| pSVA13305 | pETDuet-1-based plasmid for the overexpression of His$_6$-AviTag-*Pf*ArlI$^{T411C}$ expression in *E. coli*. | This study |
| pSVA13309 | pETDuet-1-based plasmid for the overexpression of His$_6$-AviTag-*Pf*ArlI$^{C290S/T411C}$ expression in *E. coli*. | This study |

**Table 3 - Primers**

| **Primer** | **Sequence (5' – 3')** | **Characteristics** |
|---|---|---|
| 3614 | CCCGGGGGGCCCGGGGGAAACAA CGATCTC | Forward primer for *SaarlH* down-stream containing a ApaI restriction site (underlined) |

| | | |
|---|---|---|
| 3617 | CCCCCC<u>CTGCAG</u>CCATCATAGAGGATAATGTTCC | Reverse primer for *Saarl*H up-stream containing a PstI restriction site (underlined) |
| 5167 | GATTACTACG**c**AGCAGACTACTAAAGATTACTTG | Forward primer to create the mutation *arlH*$^{E57Q}$ in pSVA2126; modified nucleotide in lower-case and bold |
| 5168 | GTAATCTTTAGTAGTCTGCT**g**CGTAGTAATCACATATCCC | Reverse primer to create the mutation *arlH*$^{E57Q}$ in pSVA2126; modified nucleotide in lower-case and bold |
| 3653 | GTGATTACTACGG**c**GCAGACTACTAAAG | Forward primer to create the mutation *arlH*$^{E57A}$ in pSVA2126; modified nucleotide in lower-case and bold |
| 3654 | CTTTAGTAGTCTGC**g**CCGTAGTAATCAC | Reverse primer to create the mutation *arlH*$^{E57A}$ in pSVA2126; modified nucleotide in lower-case and bold |
| 3659 | GTGATCACCACGG**c**ACAGACCACGAAAG | Forward primer to create the mutation *arlH*$^{E57A}$ in pSVA2100; modified nucleotide in lower-case and bold |
| 3660 | CTTTCGTGGTCTGT**g**CCGTGGTGATCAC | Reverse primer to create the mutation *arlH*$^{E57A}$ in pSVA2100; modified nucleotide in lower-case and bold |
| 5165 | CCACG**c**AACAGACCACGAAAG | Forward primer to create the mutation *arlH*$^{E57Q}$ in pSVA2100; modified nucleotide in lower-case and bold |
| 5166 | GTCTGTT**g**CGTGGTGATC | Reverse primer to create the mutation *arlH*$^{E57Q}$ in pSVA2100; modified nucleotide in lower-case and bold |
| 10765 | CAAGC**gc**ATATACTACAGTTGAATATGTTAAGC | Forward primer to create the mutation *arlH*$^{Q63A}$ in pSVA2167; modified nucleotides in lower-case and bold |
| 10766 | GTATAT**gc**GCTTGAAACATAGCTTGCGG | Reverse primer to create the mutation *arlH*$^{Q63A}$ in pSVA2167; modified nucleotide in lower-case and bold |
| 10767 | CAAGC**g**AATATACTACAGTTGAATATGTTAAGCAG | Forward primer to create the mutation *arlH*$^{Q63E}$ in pSVA2167; modified nucleotide in lower-case and bold |
| 10768 | GTATATT**c**GCTTGAAACATAGCTTGCG | Reverse primer to create the mutation *arlH*$^{Q63E}$ in pSVA2167; modified nucleotide in lower-case and bold |
| 3667 | GAAATCTAAAATC**g**CC**gc**TATCGTGGATGT | Forward primer to create the mutation *arlH*$^{T171A/S172A}$ in |

| | | |
|---|---|---|
| | | pSVA2100; modified nucleotides in lower-case and bold |
| 3668 | ACATCCACGATA**gc**GG**c**GATTTTAGATTTC | Reverse primer to create the mutation *arlH*$^{T171A/S172A}$ in pSVA2100; modified nucleotides in lower-case and bold |
| 10701 | GTGATC**gt**CACGGAACAGACCACGAAAG | Forward primer to create the mutation *arlH*$^{T55V}$ in pSVA2100; modified nucleotides in lower-case and bold |
| 10702 | TTCCGTG**ac**GATCACATAACCTTTTTTATCGCTC | Reverse primer to create the mutation *arlH*$^{T55V}$ in pSVA2100; modified nucleotides in lower-case and bold |
| 10703 | GATCACC**gt**GGAACAGACCACGAAAGATTACC | Forward primer to create the mutation *arlH*$^{T56V}$ in pSVA2100; modified nucleotides in lower-case and bold |
| 10704 | GGTCTGTTCC**ac**GGTGATCACATAACCTTT | Reverse primer to create the mutation *arlH*$^{T56V}$ in pSVA2100; modified nucleotides in lower-case and bold |
| 10778 | GTTT**gt**AGCCAATATACTACAGTTGAATATGTTAA | Forward primer to create the mutation *arlH*$^{S61C}$ in pSVA2167; modified nucleotides in lower-case and bold |
| 10779 | GTATATTGGCT**ac**AAACATAGCTTGCGGTG | Reverse primer to create the mutation *arlH*$^{S61C}$ in pSVA2167; modified nucleotides in lower-case and bold |
| 10780 | TTCA**t**GCCAATATACTACAGTTGAATATGT | Forward primer to create the mutation *arlH*$^{S62C}$ in pSVA2167; modified nucleotide in lower-case and bold |
| 10781 | ATATTGGC**a**TGAAACATAGCTTGCG | Reverse primer to create the mutation *arlH*$^{S62C}$ in pSVA2167; modified nucleotide in lower-case and bold |
| 10114 | GCAGGTTATTGGGAGAGCCCAGGTTGTGGGAGC | Forward primer to create the mutation *arlH*$^{S109}$ in pSVA2167 |
| 10115 | **a**TAGGAACCTTCTTTCCTCACTAACACCGCTAAGTAAAGGG | Reverse primer to create the mutation *arlH*$^{S109C}$ in pSVA2167; modified nucleotide in lower-case and bold |
| 10759 | <u>GAAGATTGAATGGCATGAAGGGTCC</u>ATGGCGGAAGTTATGTCACA | Forward primer to add an AviTag and His6 tag upstream of *arlI* in pSVA3116. The inserted sequence in underlined. |
| 10760 | <u>GGACCCTTCATGCCATTCAATCTTC</u>GTGGTGATGATGGTGATGGC | Reverse primer to add an AviTag and His6 tag upstream of *arlI* in pSVA3116. The inserted sequence in underlined. |
| 10761 | GGCCTGAACGACATCTTCGAAGCTCAGAAGATTGAATGGCATGAA | Forward primer to add an AviTag and His6 tag upstream of *arlI* in |

| | | |
|---|---|---|
| | | pSVA3116. The inserted sequence in underlined. |
| 10762 | TGAGCTTCGAAGATGTCGTTCAGGC**c**GTGGTGATGATGGTGATG | Reverse primer to add an AviTag and His6 tag upstream of *arlI* in pSVA3116. The inserted sequence in underlined. |
| 10774 | CCAATA**tg**CTTCATAGACAACCTCAACATA | Forward primer to create the mutation *arlI*$^{T411C}$ in pSVA5500; modified nucleotides in lower-case and bold |
| 10775 | GTCTATGAAG**ca**TATTGGGACGTTTATAGG | Reverse primer to create the mutation *arlI*$^{T411C}$ in pSVA5500; modified nucleotides in lower-case and bold |
| 10782 | ATTCGTAT**c**TGGTGAGACTGCATCTGGTAA | Forward primer to create the mutation *arlI*$^{C290S}$ in pSVA13305; modified nucleotides in lower-case and bold |
| 10783 | CTCACCA**g**ATACGAATATACTCATTCCATATTCA | Reverse primer to create the mutation *arlI*$^{C290S}$ in pSVA13305; modified nucleotides in lower-case and bold |

**Supplementary Figures 1**

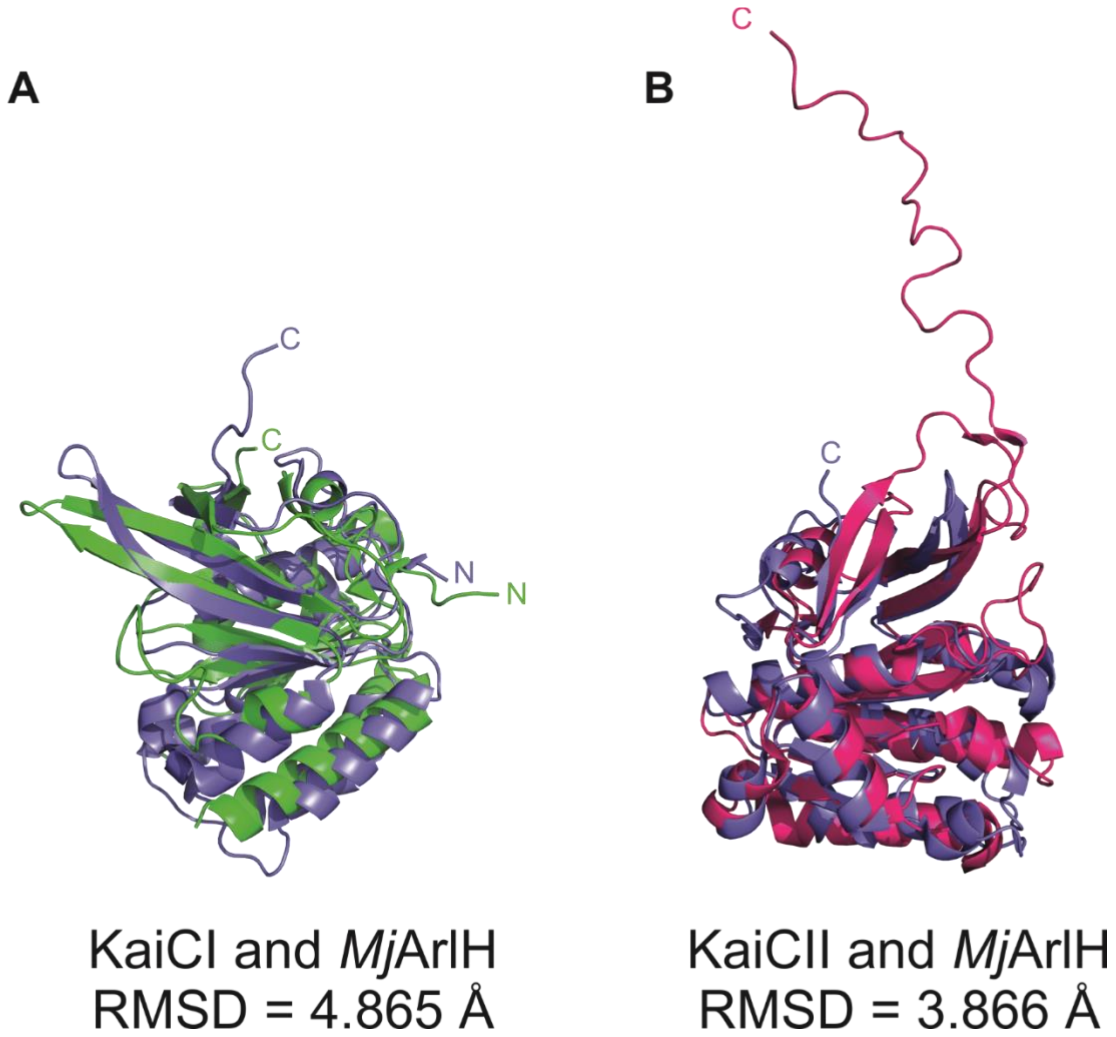


**Supp. Fig. 1** ***ArlH from S. acidocaldarius and from M. jannaschii is homologous to the KaiCII domain of KaiC***. Structural alignment **A** of 669 atoms of *Mj*ArlH (in purple; PDB: 4WIA) and KaiCI (in green; residues 16-246; PDB: 4TL8) or **B** structural alignment of 1002 atoms of *Mj*ArlH and KaiCII (in pink; residues 262-519; PDB: 2GBL). For both alignments, solvent molecules, ions, and other ligands were removed prior to structural alignment. The alignment was performed on PyMol with the command "align", and outlier atoms were removed.

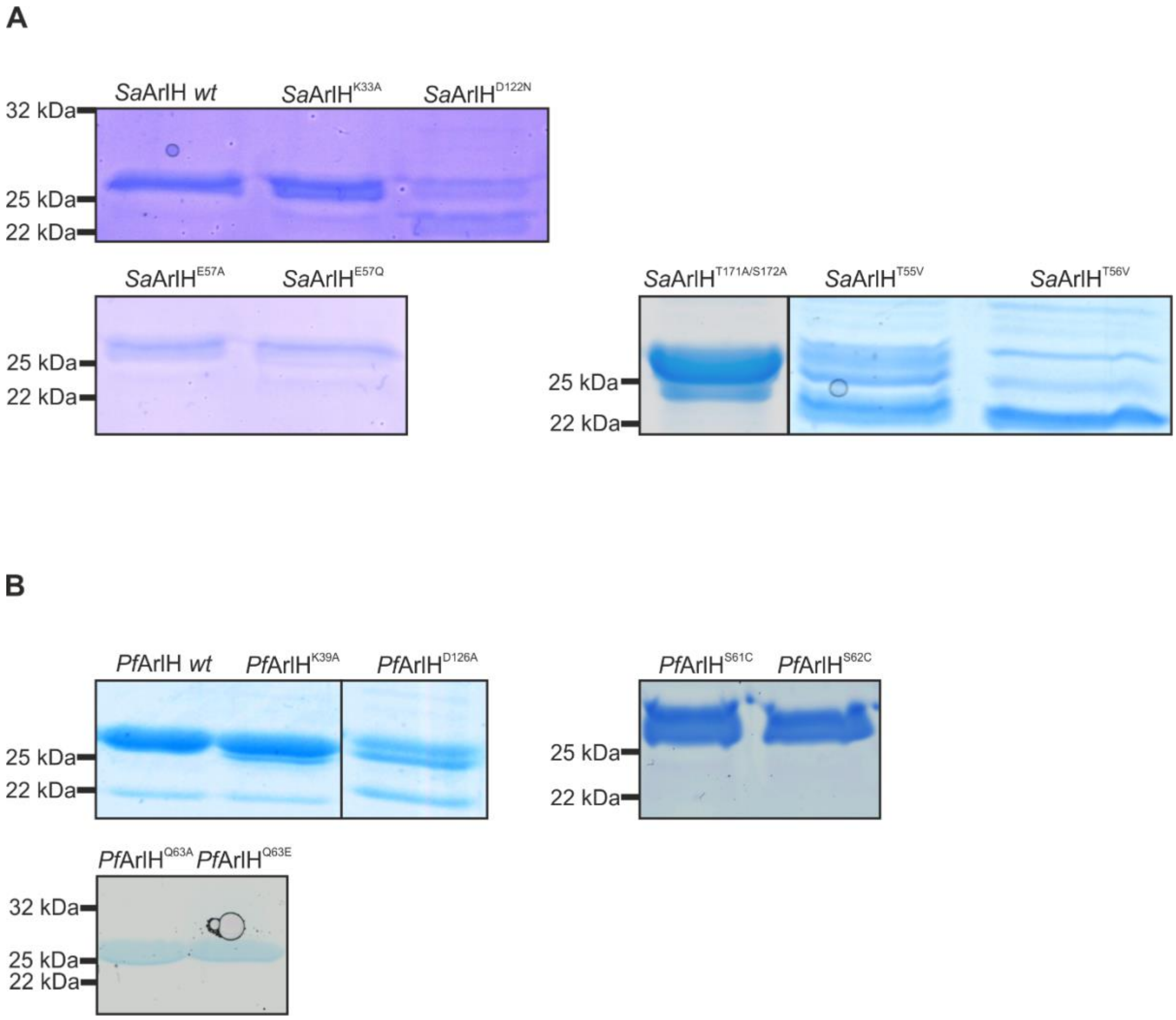


**Supp. Fig. 2** ***SDS-PAGE of purified ArlH variants*****.** ArlH from **A** *S. acidocaldarius* and **B** *P. furiosus* were purified by affinity chromatography. The Walker B mutants of both *Sa*ArlH and *Pf*ArlH are unstable, and so are the T55 mutants of *Sa*ArlH, although it was still possible to use the protein to perform MANT-ATP binding assays as indicated in Supp. Fig. 5.

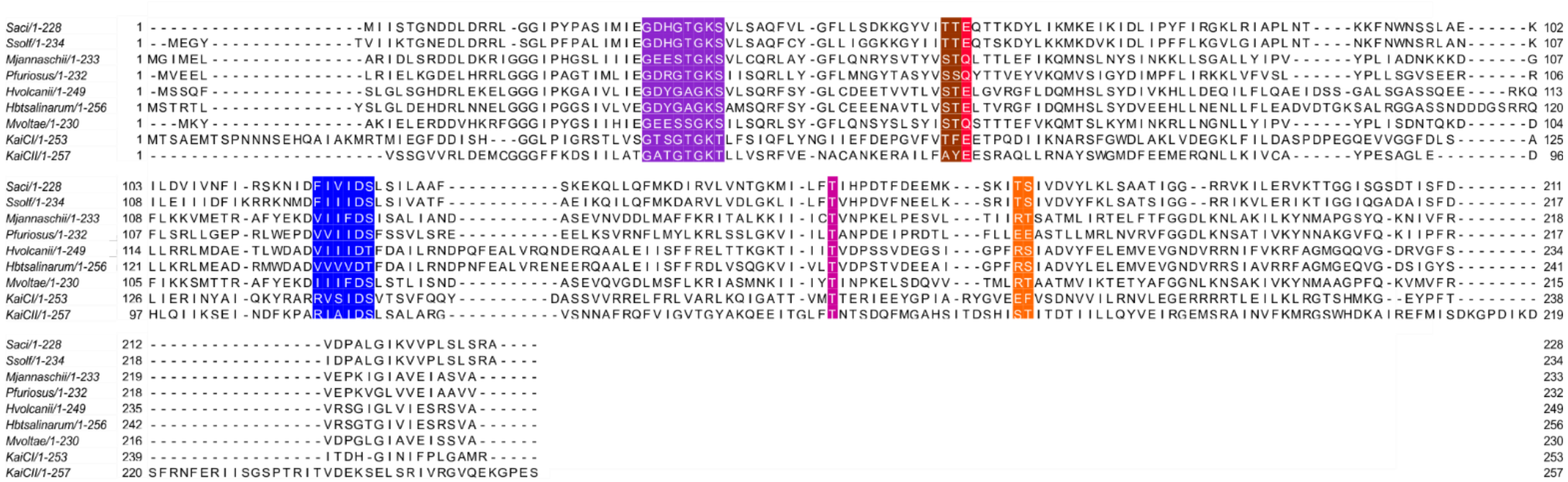


**Supp. Fig. 3** ***Alignment of different KaiC-like proteins.*** Alignment of ArlH from different organisms, indicated on the left, alongside KaiCI and KaiCII domains from *Synechococcus* (WP_011242648.1). In purple, Walker A motif; in brown, two consecutive [T/S][T/S] residues conserved in ArlH; in red, putative catalytic glutamate/glutamine; in blue, Walker B motif; in pink, T residue conserved across ArlH; in orange, phosphorylated residues in KaiCII.

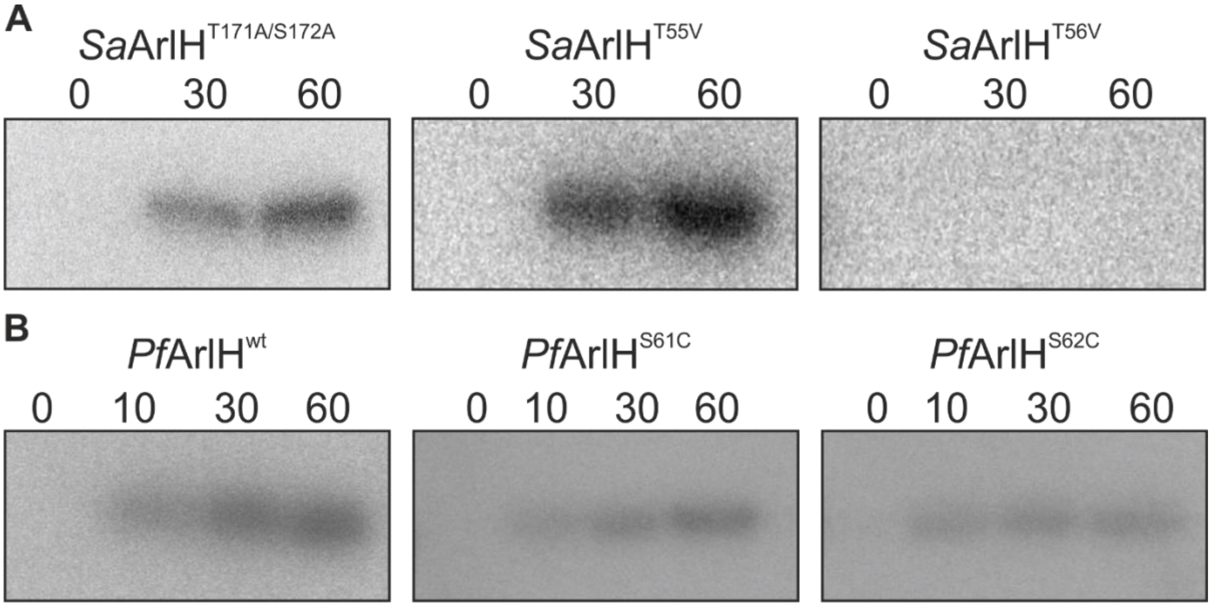


**Supp. Fig. 4** ***Conserved pairs of serine/threonine were mutated and assayed for auto-phosphorylation activity.*** **A)** phosphorylated in KaiCII. Mutating these residues to alanine does not abolish the phosphorylation signal (compare with the wild-type protein in Figure 2), indicating they are not involved in phosphorylation in *Sa*ArlH. The conserved T55 and T56 or **B**, in *P. furiosus*, S61 and S62, (Supplementary Figure 2) were hypothesised to be the phosphorylation sites. The mutation T56V abolishes phosphorylation. These mutant does not bind ATP either (Supplementary Figure 3), therefore rendering this assay inadequate to definitely settle whether this residue is phosphorylated in *Sa*ArlH. In *P. furiosus* both mutants show phosphorylation signal, suggesting that these are not the phosphorylation sites.

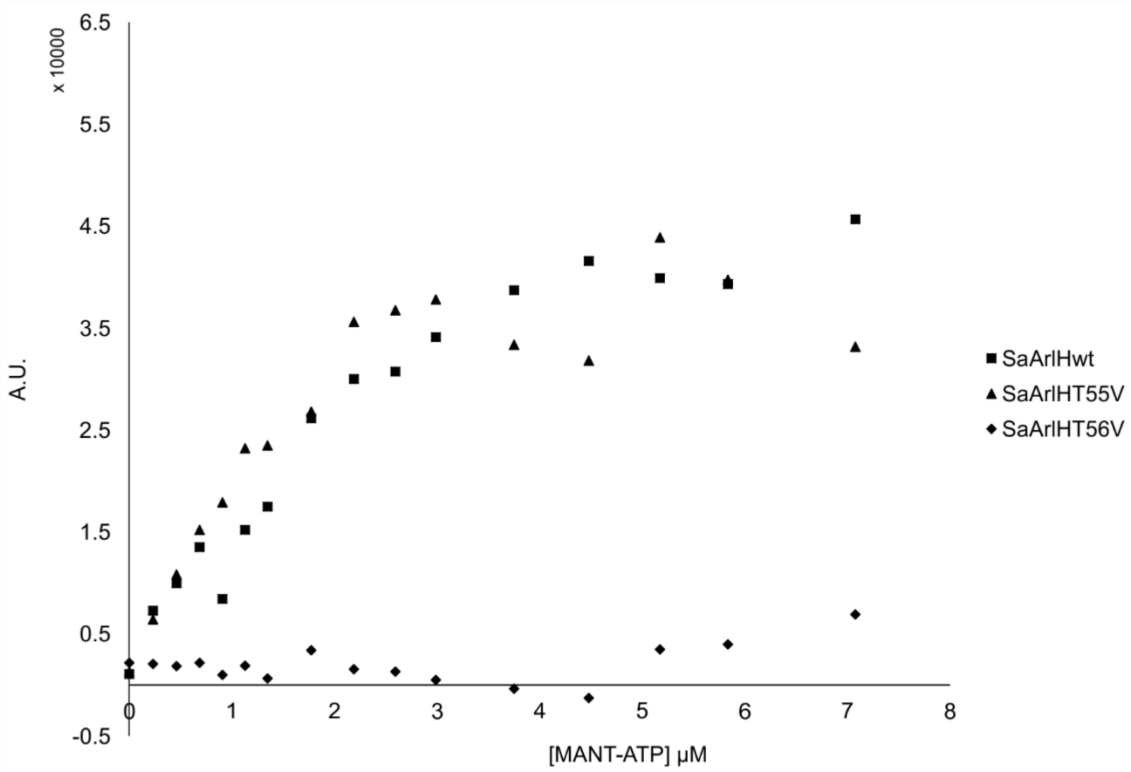


**Supp. Figure 5 Sa*ArlH*$^{T56V}$ *mutant does not bind ATP.*** Representative MANT-ATP binding assay performed with three variants of *Sa*ArlH: wild-type (squares), T55V (triangles), and T56V (diamonds). Both the wild-type and the T55V mutant can bind ATP, but this activity is severely reduced in the T56V mutant.

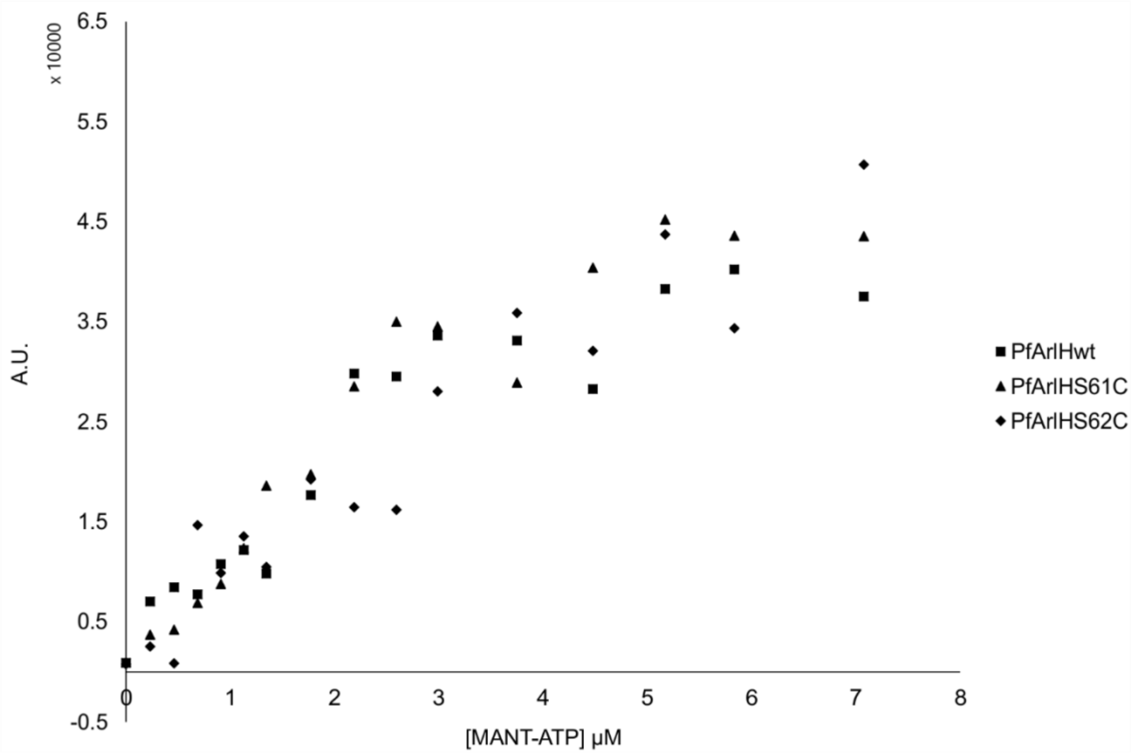


**Supp. Figure 6** ***Mutating the hypothetical phosphorylation sites of*** **Pf*ArlH does not prevent ATP-binding.*** Representative MANT-ATP binding assay performed with three variants of *Pf*ArlH: wild-type (squares), S61C (triangles), and S61C (diamonds). All three variants bind ATP with the same affinity.
